# Characterization of the upper epithelial compartment of cervical HSIL

**DOI:** 10.64898/2026.06.29.735311

**Authors:** Julia Vicari, Anna Sternjakob, Julia Staiger, Christina Kuhn, Marta Pódgorska, Justine Weishaar, Michael Döring, Tetiana Khrystenko, Yoo-Jin Kim, Erich-Franz Solomayer, Sigrun Smola

**Author notes:** Corresponding author **Correspondence** Sigrun Smola, Institute of Virology, Saarland University Medical Center, Kirrbergerstrasse, Building 47, D-66421 Homburg/Saar, Germany. Shared first authorship (Julia Vicari, Anna Sternjakob, Julia Staiger).

## Abstract

High-grade squamous intraepithelial lesions (HSIL) of the uterine cervix result from transformation by high-risk human papillomavirus (HR-HPV) and are considered precancerous lesions. Depending on the proportion of the epithelium occupied by dysplastic cells, commonly evidenced by a block-like p16INK4a staining pattern, HSIL are subdivided into CIN2 or CIN3. We used immunohistochemistry (IHC) and laser-capture microdissection (LCM) to further characterize these lesions. Broad suprabasal expression of S100A9, a component of the epidermal differentiation complex (EDC), which we had previously found to be up-regulated in genus-beta-HPV-positive skin lesions, was present in the normal exocervix and declined gradually with increasing CIN grade. In CIN2 in particular, the S100A9 staining pattern was nearly complementary to the block-like p16INK4a expression. However, it largely coincided with the expression of CD63, a marker showing strong expression in cervical crypts and in columnar cells within the normal transformation zone, also those overlying reserve cells. We then used LCM to analyze both the p16INK4a- and S100A9/CD63-positive compartments in HSIL separately. Notably, in all samples investigated, we detected the same HR-HPV type (either HPV16, 45 or 51) in both compartments of the same HSIL, while tissue samples dissected from normal exocervix or transformation zone were consistently HR-HPV-negative. Our data suggest that the upper, S100A9-positive maturing compartment of HSILs retaining weak CD63 expression may represent a residual, HR-HPV-infected columnar-cell-derived epithelium, rather than not-yet-transformed squamous epithelium that is progressively being replaced by dysplastic cells. The two-compartment architecture of these HSILs was reminiscent of the staining pattern in the normal transformation zone, where columnar cells overlie reserve cells, and compatible with HR-HPV infection of columnar-derived cells but transformation of reserve cells, the putative origin of cervical squamous cell carcinoma. Our findings have direct clinical implications. Since the upper and lower layers of HSIL appear to form distinct epithelial compartments, our study suggests that the extent and thickness of the HR-HPV-transformed lower compartment itself, rather than the proportion of the epithelium occupied by dysplastic layers, may be the decisive factor in determining the HSIL grade.

## Introduction

Cervical squamous cell carcinoma (CSCC) arises almost exclusively from persistent, transforming infection with high-risk human papillomavirus (HR-HPV) through precancerous high-grade squamous intraepithelial lesions (HSIL) (1), preferentially in the cervical transformation zone. Classical models of cervical carcinogenesis have proposed a gradual progression from cervical intraepithelial neoplasia grade 1 (CIN1) through CIN2 to CIN3, with the latter two grades together defining HSIL (2). Two models of the cellular origin of HSIL have challenged this view of gradual progression: (1) the reserve-cell model, which proposes that subcolumnar reserve cells beneath the endocervical columnar epithelium are the HR-HPV target cells and the progenitors of HSIL (3, 4); and (2) an alternative squamocolumnar-junction (SCJ) cell model, which proposed a discrete cuboidal cell population with a unique transcriptome, including CD63, as the cancer-prone cell (5). HSIL is characterized by deregulated viral E6/E7 expression, which is reflected by block-like p16INK4a-positivity of the lesion (6). Recent approaches that further separate productive from transforming lesions include staining for the HPV E4 protein (7). Within HSIL, the distinction between CIN2 and CIN3 relies on the proportion of the epithelium occupied by dysplastic cells with an increased nuclear-to-cytoplasmic ratio, nuclear pleomorphism and mitoses. In CIN3, dysplastic cells occupy more than two-thirds and up to the full thickness of the squamous epithelium, with little or no surface layer remaining; in CIN2, dysplastic cells occupy up to the lower two-thirds. Although CIN2 has lower diagnostic reproducibility than CIN3 (8) the distinction carries clinical weight, because the management of the less progressed CIN2 may be more conservative, particularly in younger women, according to current guidelines (9).

We hypothesized that S100A9 (calgranulin B), an EDC protein, which we had previously investigated in the context of cutaneous genus beta-HPV lesions (10) could shed more light into the nature of the upper non-transformed HSIL compartment. We therefore investigated its expression in different CIN lesions and in normal human cervix.

### Materials and methods Immunohistochemistry (IHC)

Anonymized FFPE cervical tissue specimens were retrieved from the archives of the Department of Pathology, Saarland University Medical Center, Germany. The initial diagnosis and grading had been established by expert pathologists. Sections (1-4 µm) were cut for IHC. After epitope retrieval in 10 mM sodium citrate (pH 7.0), the slides were blocked with 2.5% normal horse serum (Vector, Burlingame, CA, USA) and incubated with antibodies (Abs) against p16INK4a (1:64; CINtec, Roche/Ventana, #9511), S100A9 (1:4000; Santa Cruz, #sc-20173, clone H-90), CD63 (1:2000; Thermo Scientific, #MA5-11501) and p63 (1:200; Abcam, #ab735, clone 4A4). Detection was performed with the ImmPRESS peroxidase (anti-mouse; Vector, #MP-7402) or ImmPRESS AP (anti-rabbit; Vector, #MP-5401-15) reagents, followed by hematoxylin counterstaining. Slides were evaluated on a Leica DMI 6000B microscope (Leica, Wetzlar, Germany).

### Laser-capture microdissection (LCM) and HPV detection

FFPE sections of CIN2 and CIN3 lesions were used for LCM. To avoid HPV-DNA carry-over the microtome was cleaned using RNase AWAY® (Molecular Bio-Products, Inc., San Diego, CA, USA) between cases and the microtome blade was changed after each cut according to (11, 12). Sections (10 µm) were mounted onto UV-irradiated, membrane-coated slides suitable for LCM (Zeiss, #4151909081000). Slides were dried at 56 °C overnight. To allow morphological identification of the epithelial lesions, sections were stained with a rapid hematoxylin protocol following the manufacturer’s recommendations for the Zeiss PALM MicroBeam system used for LCM. Regions of interest were identified by comparison with adjacent p16INK4a- and S100A9-stained IHC slides. Target areas were selectively dissected (Supporting Figure 1) and catapulted into adhesive caps 500 (Zeiss, #4151909201000) using UV laser pulses under standard instrument settings (PALMRobo 4.9 Pro software).

### DNA extraction, HPV PCR and genotyping

Genomic DNA was extracted from microdissected FFPE-material with the QIAamp DNA Micro Kit (Qiagen, #56304), following the Zeiss PALM application guidelines. Briefly, the collected tissue fragments were lysed overnight in the cap in proteinase K-containing buffer at 56 °C, and DNA was purified on spin columns according to the manufacturer’s instructions. DNA was eluted in 20 µl distilled water and genotyped with INNO-LiPA HPV Genotyping Extra II (Fujirebio Germany GmbH, Hannover, Germany).

## Results

### Gradual loss of suprabasal S100A9 expression in cervical intraepithelial neoplasia

We examined S100A9 expression in normal human exocervix and in cervical intraepithelial neoplasia grades 1-3. In normal exocervical epithelium, S100A9 showed strong suprabasal expression with a negative basal and adjacent suprabasal layer (Figure 1A), consistent with its role as a squamous differentiation factor (13). In CIN1 and CIN2, S100A9 staining remained confined to the upper layers but spared an increasing proportion of suprabasal cells (Figure 1B, C). In CIN3 the S100A9 staining was either absent, in some CIN3 it was detected only in the most superficial layers or it showed a more patchy pattern as seen in the lower part of CIN in Figure 1C. Infiltrating cells with strong S100A9 positivity, typical of myeloid cells, were also observed confirming proper IHC staining in CIN3 with almost completely negative epithelium (Figure 1D).

**Figure 1.**
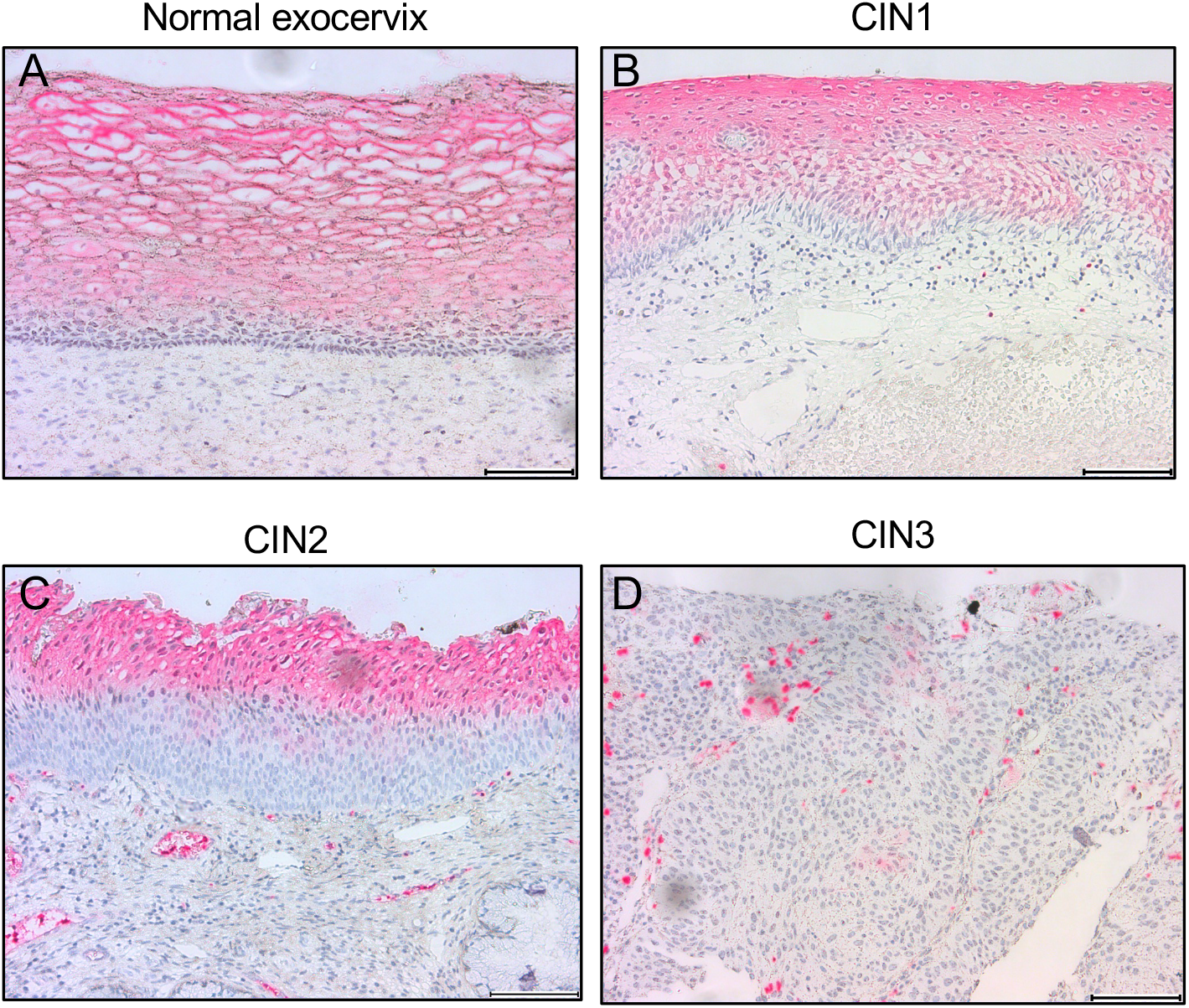
Suprabasal S100A9 expression in exocervix and decline from CIN1 to CIN3. Human FFPE-sections were stained with anti-S100A9 Ab (red color) and counterstained with hematoxylin. (A) Representative sections of exocervix (n=10), (B) CIN1 (n=7), (C) CIN2 (n=8), (D) CIN3 (n=15); scale bars: 100 µm.

Overall, there was a gradual loss of S100A9 positivity as the proportion of dysplastic layers increased, consistent with the view that the differentiated squamous compartment is progressively replaced from below by dysplastic cells during the development of cervical precancer.

### S100A9 is expressed in the compartment above the block-like p16INK4a staining but can extend laterally beyond it

Evaluation of the same CIN2 lesion as in Figure 1C at lower magnification showed that S100A9 was expressed across a broad stretch of the upper part of the epithelium (Figure 2A), located above the block-positive part of the p16INK4a staining (Figure 2B), indicating an almost complementary distribution of the two markers. Laterally, however, the two-part architecture, with S100A9-positivity above, extended well beyond the boundary of the underlying p16INK4a-positive area. This indicated that HR-HPV transformation, as evidenced by p16INK4a positivity, is not a prerequisite for the apparent two-compartment architecture of CIN2.

**Figure 2.**
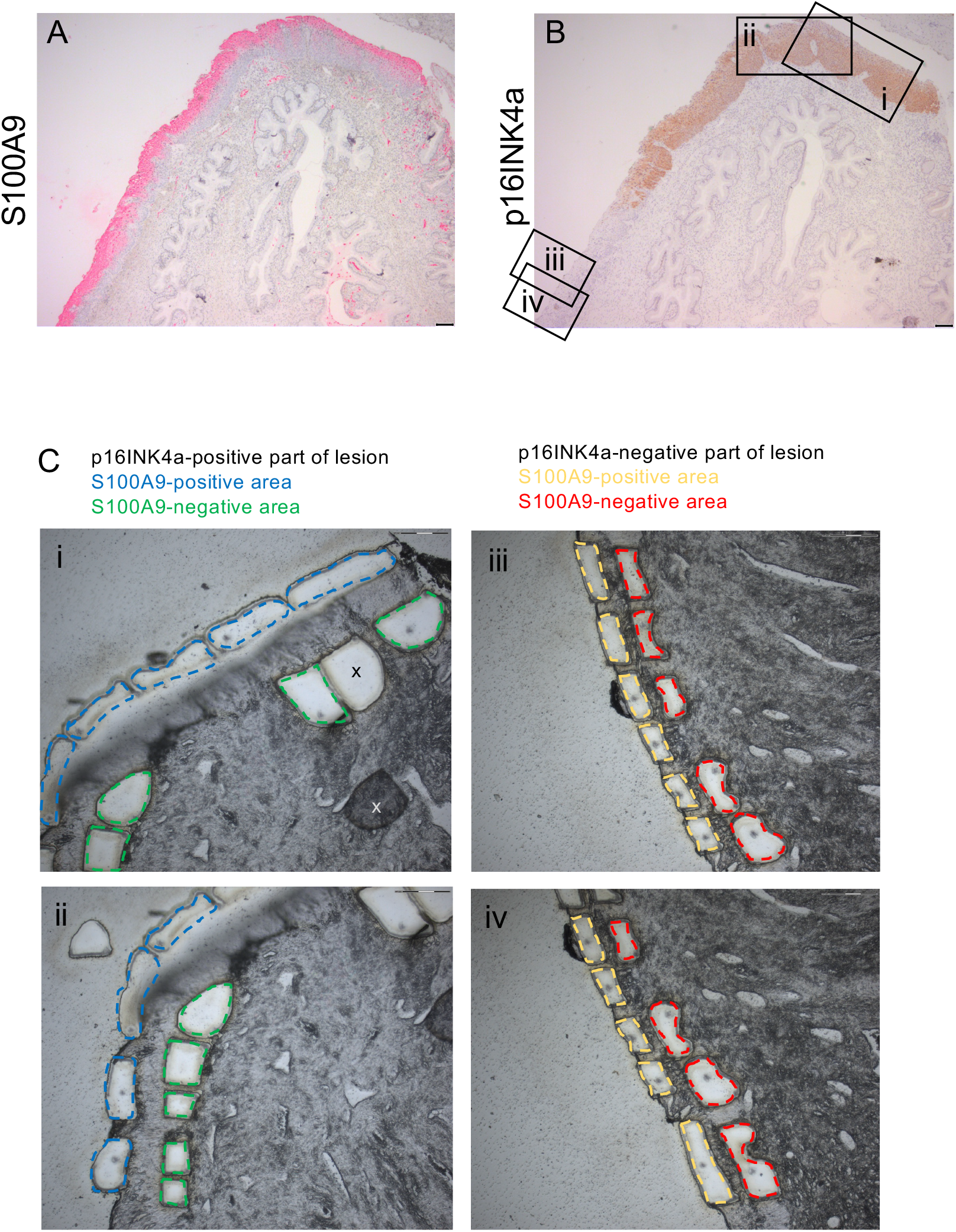
S100A9, p16INK4a staining pattern and LCM-based genotyping of the upper and lower CIN2 compartments. Human HSIL FFPE-sections (n=6) were stained with (A) anti-S100A9 Ab (red color), (B) anti-p16INK4a Ab (brown color) and counterstained with hematoxylin; scale bars: 100 µm. A representative section of CIN2 (identical lesion as in Figure 1C) is shown; (C) LCM of FFPE-sections from HSIL (n=5). Shown is the same CIN2 lesion as in A and B. Dissected areas that were pooled (from i and ii, or from iii and iv, respectively) and used for HPV genotyping are marked with dashed lines. Left panel: upper S100A9-positive area (dashed blue line), lower S100A9-negative but p16INK4a-block-positive area (dashed green line); right panel: upper S100A9-positive area (dashed yellow line), lower S100A9-negative and p16INK4a-block-negative area (dashed red line); x: LCM-cut tissue piece not properly catapulted; scale bars: 150 µm.

To determine the HR-HPV DNA status of each compartment, the upper (S100A9-positive) and lower (p16INK4a block-positive or -negative) layers of CIN2 (n=4) and one CIN3 lesion were carefully separated from one another, excised using the LCM method, leaving a non-dissected part in between (Figure 2C), and analyzed for HR-HPV DNA. Two CIN2 and the CIN3 lesion were positive for HR-HPV16, and one each for HR-HPV45 or HR-HPV51. The CIN2 lesion shown in Figures 1C, 2 and 4A was positive for HR-HPV51 in the p16INK4a-positive lower part. Interestingly, we additionally detected HR-HPV51 not only in the upper, S100A9-positive layers above the p16INK4a-block staining but also in the p16INK4a-negative region beneath the remaining S100A9-positive band. Also in other microdissected HSIL, identical HR-HPV were detected in both the upper and lower compartments of the same lesion (Table 1). LCM was also performed on control tissue from normal exocervix and transformation zone (n=2). This included metaplasia and columnar cells overlying reserve cells, which proved HR-HPV negative (Table 1), while we were unable to extract DNA from the reserve cells. These findings indicate that both the upper and the lower compartment of CIN2 can harbor HR-HPV, but that only the lower compartment is transformed by HR-HPV as indicated by p16INK4a-block positivity.

**Table 1.**
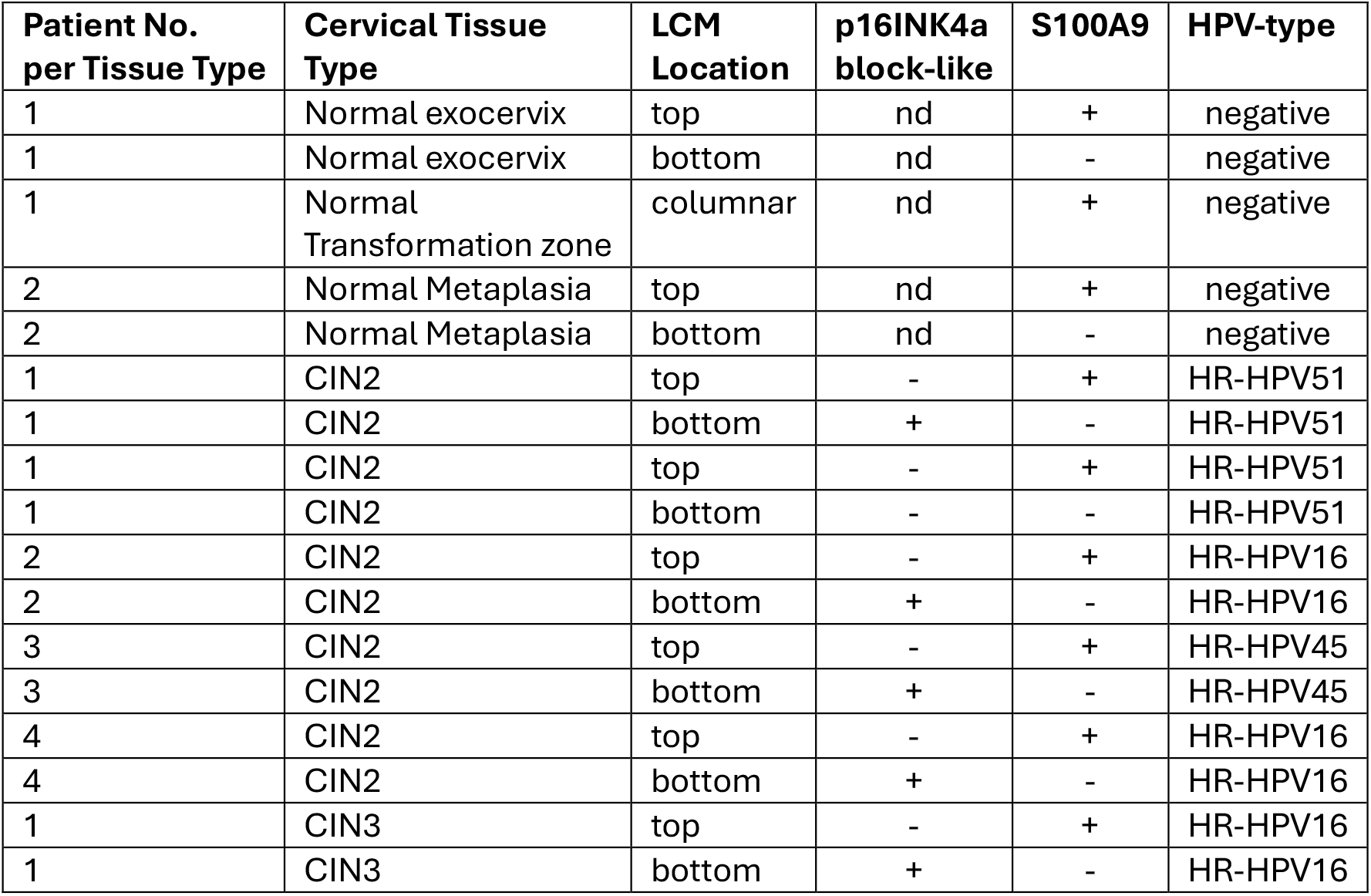
HPV-Genotyping of laser-capture microdissected cervical tissue from n=5 patients, n=2 normal cervical tissue, n=4 CIN2, n=1 CIN3. Shown are the results of HPV genotyping, p16INK4- and S100A9-IHC per (pooled) tissue from respective LCM-locations, documented in Figure 2C and Supplementary Figure 1; nd= not done.

| Patient No.<br>per Tissue Type | Cervical Tissue<br>Type | LCM<br>Location | p16INK4a<br>block-like | S100A9 | HPV-type |
| --- | --- | --- | --- | --- | --- |
| 1 | Normal exocervix | top | nd | + | negative |
| 1 | Normal exocervix | bottom | nd | - | negative |
| 1 | Normal<br>Transformation zone | columnar | nd | + | negative |
| 2 | Normal Metaplasia | top | nd | + | negative |
| 2 | Normal Metaplasia | bottom | nd | - | negative |
| 1 | CIN2 | top | - | + | HR-HPV51 |
| 1 | CIN2 | bottom | + | - | HR-HPV51 |
| 1 | CIN2 | top | - | + | HR-HPV51 |
| 1 | CIN2 | bottom | - | - | HR-HPV51 |
| 2 | CIN2 | top | - | + | HR-HPV16 |
| 2 | CIN2 | bottom | + | - | HR-HPV16 |
| 3 | CIN2 | top | - | + | HR-HPV45 |
| 3 | CIN2 | bottom | + | - | HR-HPV45 |
| 4 | CIN2 | top | - | + | HR-HPV16 |
| 4 | CIN2 | bottom | + | - | HR-HPV16 |
| 1 | CIN3 | top | - | + | HR-HPV16 |
| 1 | CIN3 | bottom | + | - | HR-HPV16 |

### S100A9 and CD63 expression patterns in columnar epithelial cells of the normal human transformation zone

Since the two compartments (upper S100A9-positive and lower S100A9-negative) appeared to exist prior to HR-HPV transformation, we examined the normal transformation zone, where HSIL originates.

Consistent with its nature as a squamous EDC marker, we detected S100A9 expression in maturing squamous metaplasia (Figure 3A), which was HPV-negative in the S100A9-positive as well the -negative parts (Table 1). Notably, S100A9 was expressed even more prominently in columnar epithelial cells (Figure 3B), including those overlying p63-positive reserve cells (Figure 3B, C). Thus, S100A9 expression in the cervical epithelium was not restricted to differentiating squamous keratinocytes but extended to columnar cells. Infiltrating cells with strong S100A9 positivity, typical of myeloid cells, were also observed.

**Figure 3.**
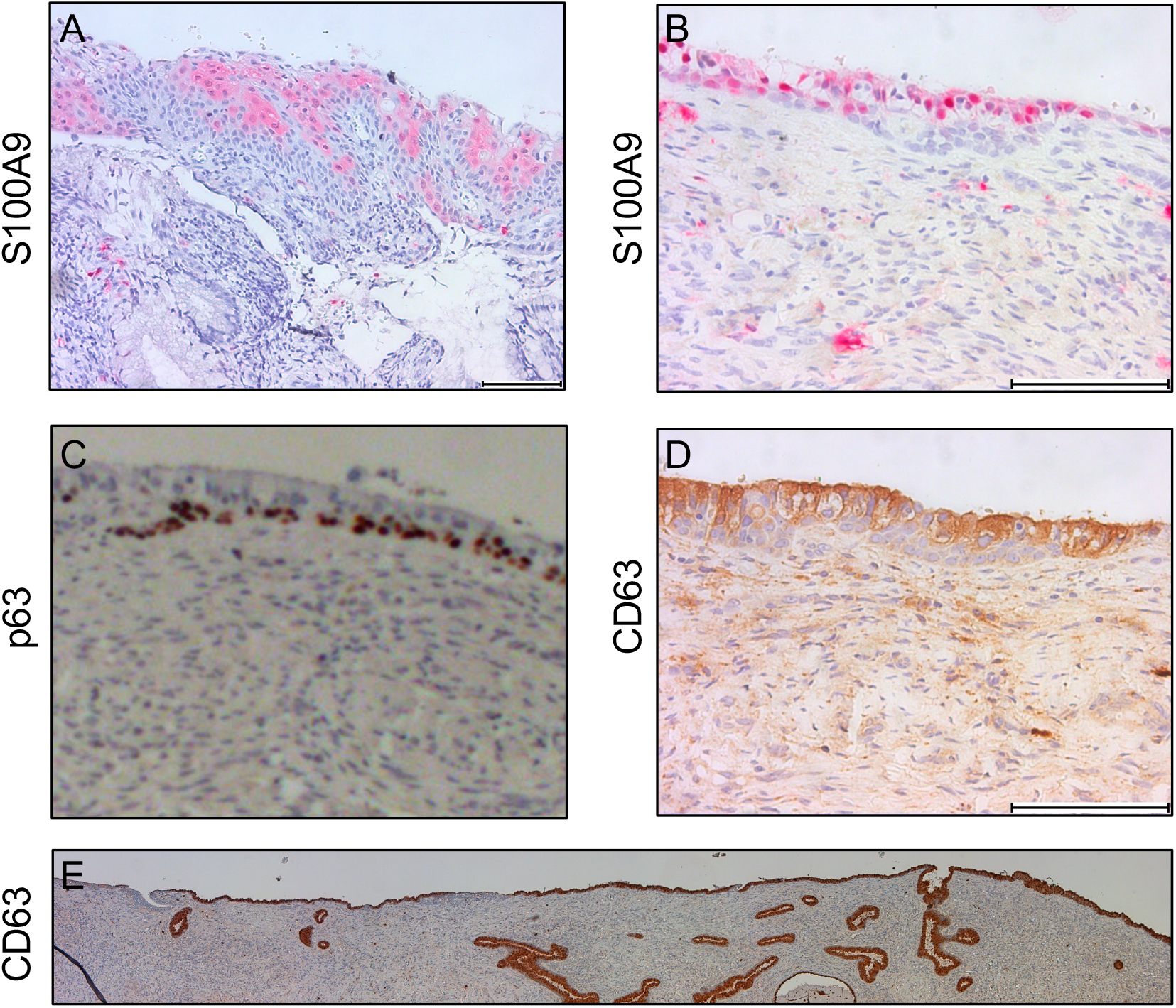
S100A9 and CD63 expression patterns in normal transformation zone. FFPE-sections of the human normal cervical transformation zone were stained with anti-S100A9 Ab (A and B, n=2), in (A) maturing metaplasia and in (B) columnar epithelium in (red color), with anti-p63 Ab (brown color) to identify reserve cells (C), with anti-CD63 Ab (brown color) (D and E), and counterstained with hematoxylin; scale bars: 100 µm. (B-D) Consecutive sections of the same representative transformation zone. (E) Transformation zone of an independent donor, overview.

We next stained normal tissue for CD63, a marker previously reported in cuboidal cells at the SCJ (5), and detected CD63 expression in columnar epithelial cells near the SCJ overlying p63-positive reserve cells (Figure 3C, D). Unexpectedly, CD63 was strongly expressed almost throughout the endocervical columnar epithelium as well as in endocervical crypts (Figure 3E).

These data indicate that (1) the columnar epithelium is able to activate a S100A9 program, as previously observed in columnar epithelia of other organs (14); (2) this includes columnarvepithelial cells overlying reserve cells; and, unexpectedly, (3) CD63 expression in the endocervical epithelium is not restricted to a small cell population at the SCJ as previously described (5).

### CD63 expression in the S100A9-positive upper layers of CIN2

On the basis of these observations in the normal transformation zone, we further characterized CIN2 with the same anti-CD63 Ab that stained columnar cells in the normal transformation zone. Notably, the upper compartment of the CIN2 lesion shown in Figures 1 and 2 showed band-like CD63 staining in the S100A9-positive compartment overlying the p16INK4a-positive area (Figure 4A). Compared with the crypts in the same tissue section, however, staining was weaker. CD63 expression in the upper HSIL compartment (Figure 4A and B) and its complementary relationship to the block-like p16INK4a staining (Figure 2B, Figure 4C showing the same HR-HPV16-positive lesion as in Figure 4B) were consistent features of HSIL, particularly of CIN2 lesions, and were independent of the HR-HPV type.

**Figure 4.**
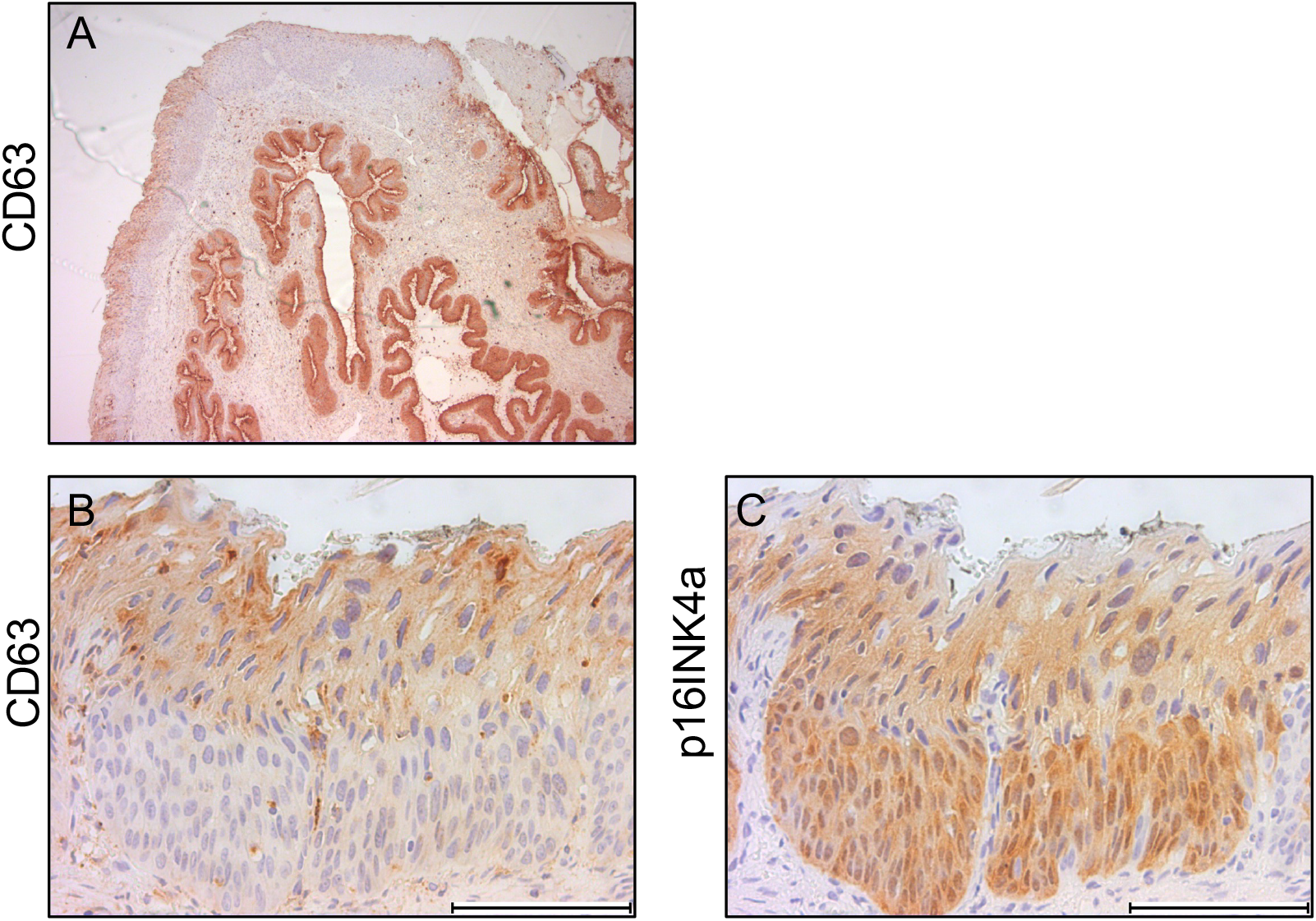
Superficial layers in CIN2 express CD63. (A) Human FFPE-section from HR-HPV51-positive CIN2 (identical lesion as in Figures 1C and 2) was stained with anti-CD63 Ab (brown color); human FFPE-sections from different HSILs (n=7) were stained with (A and B) anti-CD63 Ab (brown color), or (C) anti-p16INK4a Ab (all brown color). (B and C) Shown is a representative HR-HPV16-positive CIN2; counterstain hematoxylin; scale bars: 100 µm.

Together, these data suggest for the first time that the epithelial layers overlying the HR-HPV-transformed dysplastic lesion in HSIL express features found in columnar cells, including those overlying reserve cells.

## Discussion

In this study we identified a two-compartment architecture of HSIL, particularly in CIN2, with a distinct profile of the layers above the p16INK4a-positive compartment. The upper compartment expressed S100A9, and, at the first glance, morphologically resembled differentiating squamous epithelium that is “not yet” transformed. However, we clearly showed that it retains residual expression of the marker CD63, normally expressed in columnar endocervical cells including those overlying reserve cells. Our data provide evidence that this two-compartment architecture exists in the normal transformation zone before HR-HPV transformation, and suggest that the upper S100A9/CD63-positive HSIL compartment is likely to be derived from columnar cells.

### S100A9 expression is not restricted to squamous-derived epithelium

We found robust S100A9 expression not only in suprabasal squamous cells but also in columnar epithelial cells of the normal endocervix and in columnar cells overlying reserve cells in the transformation zone. Importantly, the presence of S100A9 in normal columnar cells means that S100A9 cannot be regarded as a pure squamous-lineage marker. In fact, expression of S100A8/A9 by columnar and glandular epithelia is not unprecedented. In the airways, bronchial columnar epithelial cells up-regulate S100A8/A9 in response to inflammatory stimuli (14), and in the porcine uterus the endometrial (columnar) epithelium expresses S100A8/A9 under hormonal and inflammatory control (15). The mechanism driving S100A9 expression in cervical columnar cells is unknown but it is plausible to assume that it relates to the microenvironment of the transformation zone. Candidate stimuli include the locally low and fluctuating pH of the transformation zone, cellular stress as well as and hormonal influences given the documented steroid hormone- and cytokine-responsiveness of S100A8/A9 in porcine endometrial epithelium (15).

### The upper compartment of CIN2 is likely columnar-derived rather than a residual squamous epithelium

The upper compartment of CIN2 has long been interpreted as a residuum of squamous maturation. Our data challenge this interpretation. The upper, S100A9-positive compartment retained CD63, a marker, which we found to be broadly expressed by endocervical columnar epithelium. The same two-layer arrangement (S100A9/CD63-positive cells overlying reserve cells) was already present in the normal transformation zone. Although, the tetraspanin CD63 may also be expressed by other cell types, the most intriguing explanation is that the upper compartment of CIN2 is a residual columnar-derived epithelium that overlies and is progressively undermined by an expanding HR-HPV-transformed population beneath it. The hypothesis that a two-compartment structure is present prior to transformation is further supported by our observation that the S100A9/CD63-positive layers extended laterally well beyond the underlying p16INK4a-block-positive epithelium. Notably, CD63 was one of the markers originally used to define the proposed “discrete” SCJ cell population (5); our observation that CD63 is broadly and strongly expressed by endocervical columnar cells including crypt cells, rather than confined to a small junctional population, is consistent with recent high-resolution mapping of the transformation zone that likewise did not identify a discrete CK7-positive junctional cell type (16).

### HR-HPV transformation is confined to the lower compartment, consistent with the reserve-cell hypothesis

Although our descriptive study cannot clarify the origin of the p16INK4a-positive lower compartment, the two-compartment architecture we describe here is consistent with the reserve cell model of cervical carcinogenesis, according to which HSIL arises from HR-HPV-infected, proliferating reserve cells located beneath the columnar epithelium, which are distributed across an extended endocervical zone (3, 4, 17). Our findings are also compatible with the recently described progression through “thin” HSIL, a reserve-cell-derived lesion that shares genomic alterations with thick HSIL and invasive carcinoma (18). The recent demonstration that deregulated HR-HPV E6/E7 transcription is concentrated in reserve cells provides independent in situ support for this model (16). Future studies will be needed to clarify the relationship between the two compartments we have described in order to clarify whether reserve cells are (1) derived from the overlying columnar cells, (2) share a common precursor with them, or (3) represent an independent, embryologically distinct progenitor population (17, 19-21).

### The upper CD63-positive compartment can be HR-HPV-infected without being transformed

Our LCM-derived PCR data showed the same HR-HPV types in both compartments of CIN2. This indicates that HR-HPV can infect the upper compartment but does not transform it, in the sense that it does not induce the block-like p16INK4a phenotype. Our findings are consistent with the report of infection of normal-appearing junctional/columnar cells by transcriptionally active HR-HPV (22), and the view that the cellular context may determine the progression and histological subtype of cancer (23). Since our genotyping data are based on microdissection of adjacent compartments, contamination must, however, be rigorously excluded; lack of cross-contamination in our study is supported by the detection of distinct HR-HPV types across the different lesions, by HR-HPV-negative normal control samples treated with LCM in the same manner and the contact-avoiding method of the LCM technique itself. HR-HPV in situ hybridization, however, would provide more definitive, spatially resolved confirmation and is a priority for future work.

### Implications for the distinction of CIN2/CIN3

The conventional grading of HSIL uses the proportion of the epithelium occupied by dysplastic cells to separate CIN2 from CIN3. If, however, the maturing upper layer is a retained, columnar-derived compartment, this raises important questions: (1) whether the presence and relative thickness of this superficial epithelium, which currently distinguishes CIN2 from CIN3, can truly account for a different biological behavior and prognosis, or (2) whether the biologically decisive variable is instead the size (extent and thickness) of the HR-HPV-transformed compartment itself, i.e. the amount of p16INK4a-positive epithelium.

## Conclusions

The upper, maturing compartment that defines CIN2 does not appear to be a “not yet transformed” squamous differentiated epithelium but could rather derive from columnar epithelium. It retains CD63 as a marker of columnar cells in the transformation zone, can express S100A9, and harbor HR-HPV. However, this layer appears to be less prone to HR-HPV transformation, since block-like p16INK4a positivity is rather confined to the compartment beneath. Notably, a two-compartment architecture is already present in the normal transformation zone, before HR-HPV transformation. These observations support the reserve-cell hypothesis of cervical cancer origin and a progression model, in which the transformed compartment expands beneath a pre-existing columnar layer, rather than the progression from productive HR-HPV-infection via CIN1 to CIN2 and 3.

Furthermore, our data suggest that the morphological criterion currently used to distinguish between CIN2 and CIN3, the proportion of the epithelium occupied by dysplastic cells, may reflect the extent to which the superficial layer remains intact rather than an intrinsic difference in the transforming lesion; the extent and thickness of the p16INK4a-positive area may be a more meaningful measure. For clinicians, this new perspective could refine the risk stratification of HSIL and thus influence decisions regarding conservative or surgical treatment, particularly in young women.

## Acknowledgments

The authors thank Katrin Thieser, Anna Werle, Barbara Best, Angelina Glahn and Anne Kerber for excellent technical assistance, and Ulrike Fischer for advice on the LCM.

## Ethics and AI statements

This retrospective study using anonymized pathology samples was approved by the institutional review board of the Saarland Ärztekammer and carried out in accordance with the principles of the Declaration of Helsinki.

During the preparation of this work, the authors used claude.ai (Anthropic) and DeepL in order to improve sentence structure and language. After using this tool, the authors reviewed and edited the content as needed and take full responsibility for the content of the published article.

## Supporting Figure 1

**Supporting Figure 1.**
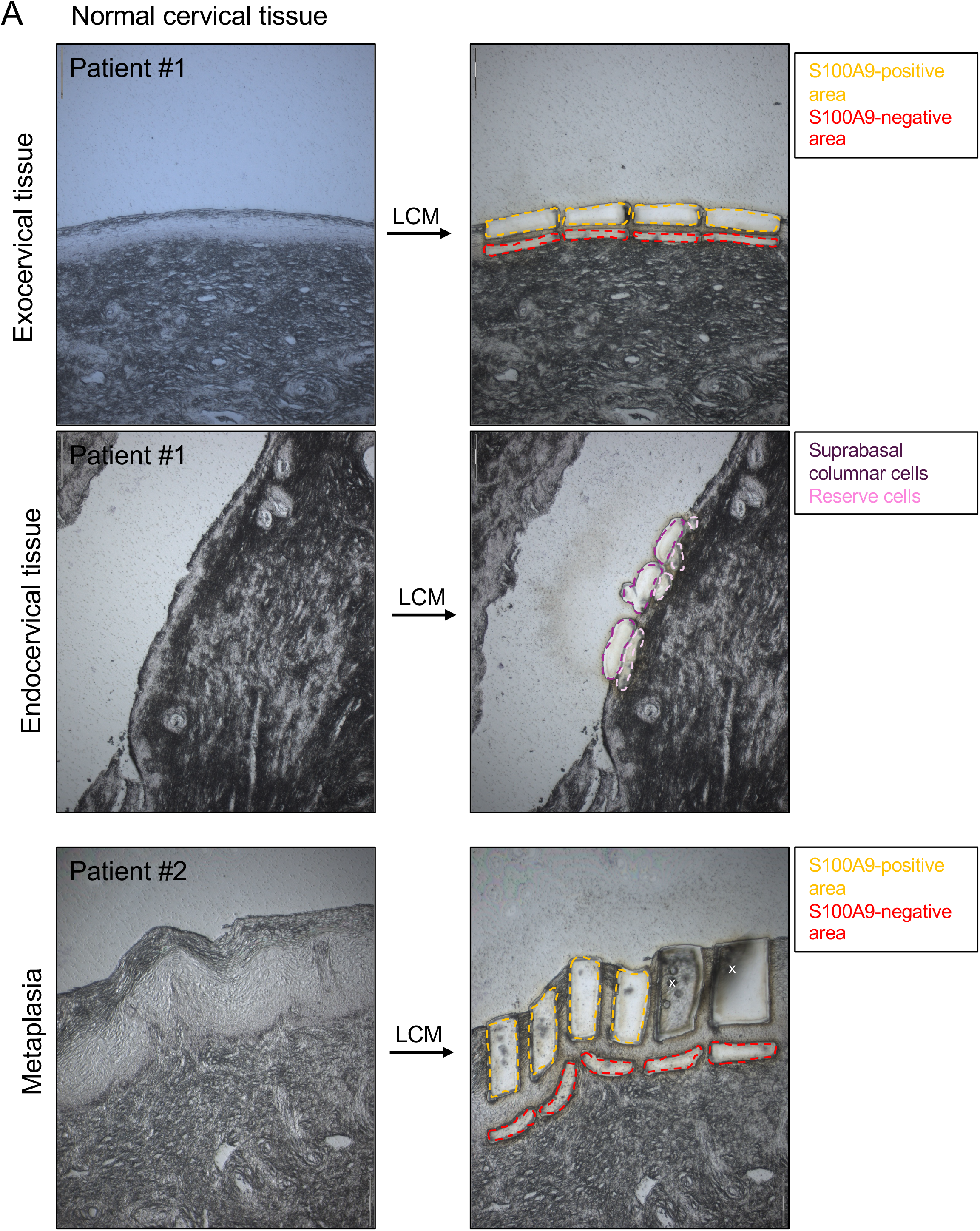
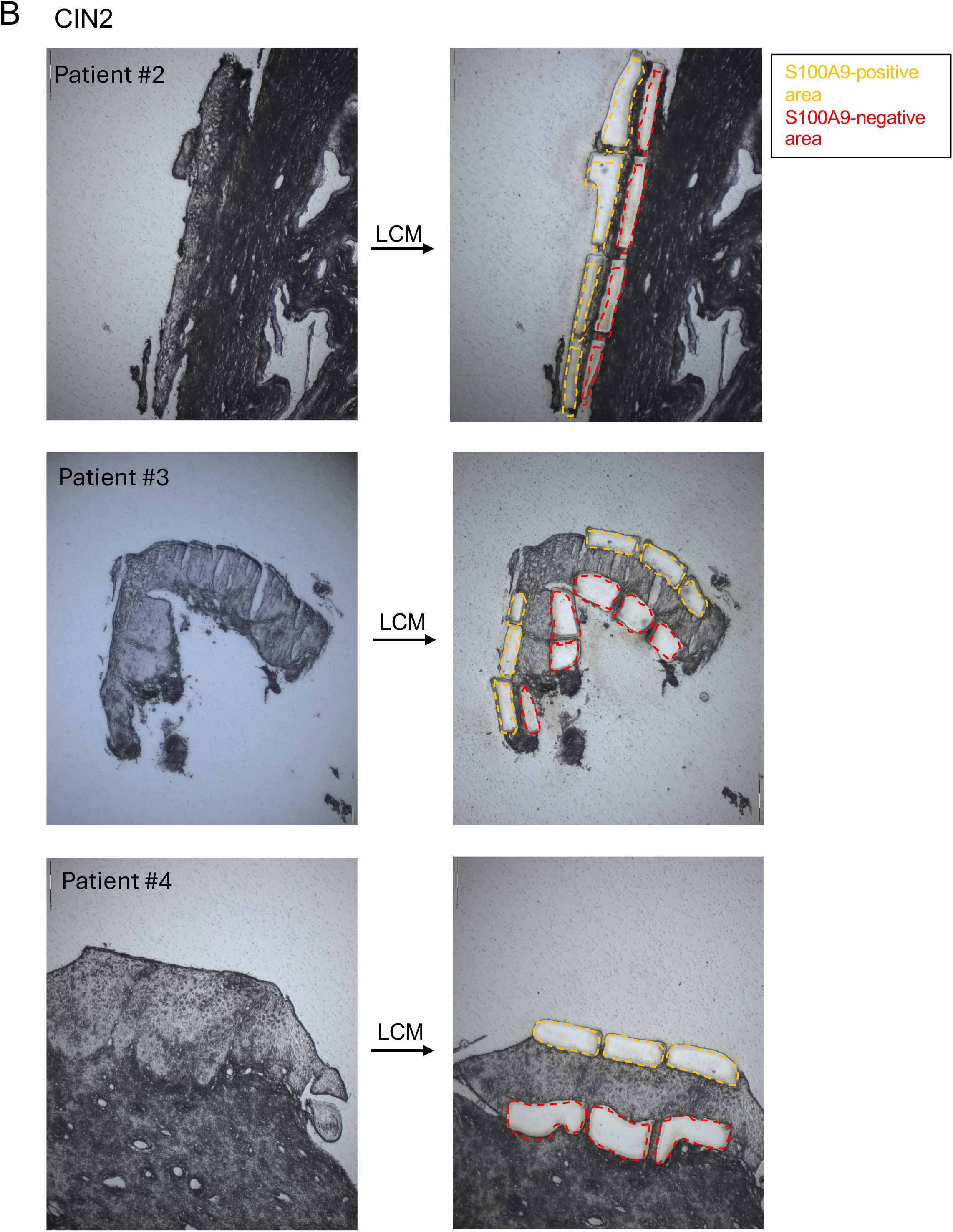
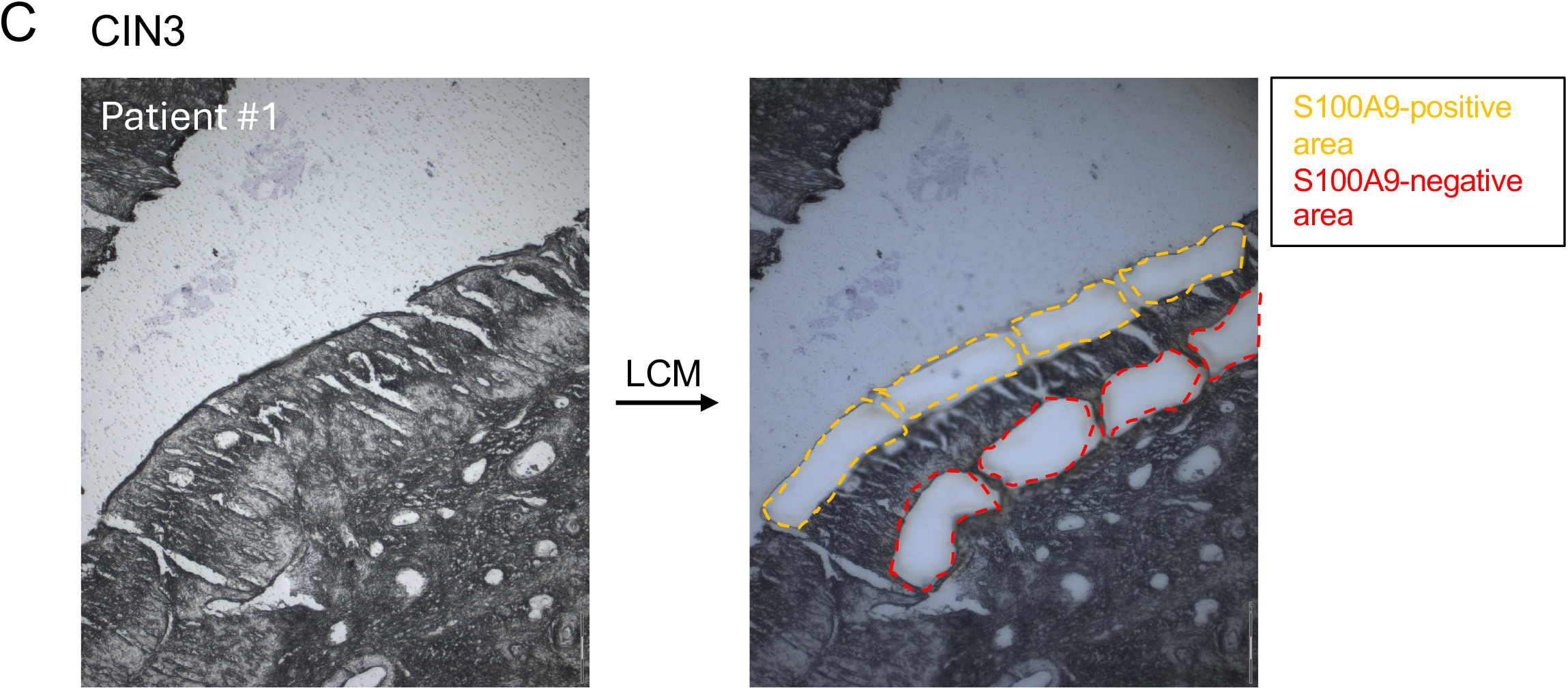
LCM of FFPE-sections from (A) normal cervix (n=2), (B) CIN2 (n=4; CIN2 from patient #1 is shown in Figure 2C) and CIN3 (n=1). Dissected areas that were pooled and used for HPV genotyping are marked with dashed lines as indicated. x: LCM-cut tissue pieces not properly catapulted; scale bars: 150 µm.

